## Supplementary Figures Merged for "Genetic Dissection of the RNA Polymerase II Transcription Cycle"

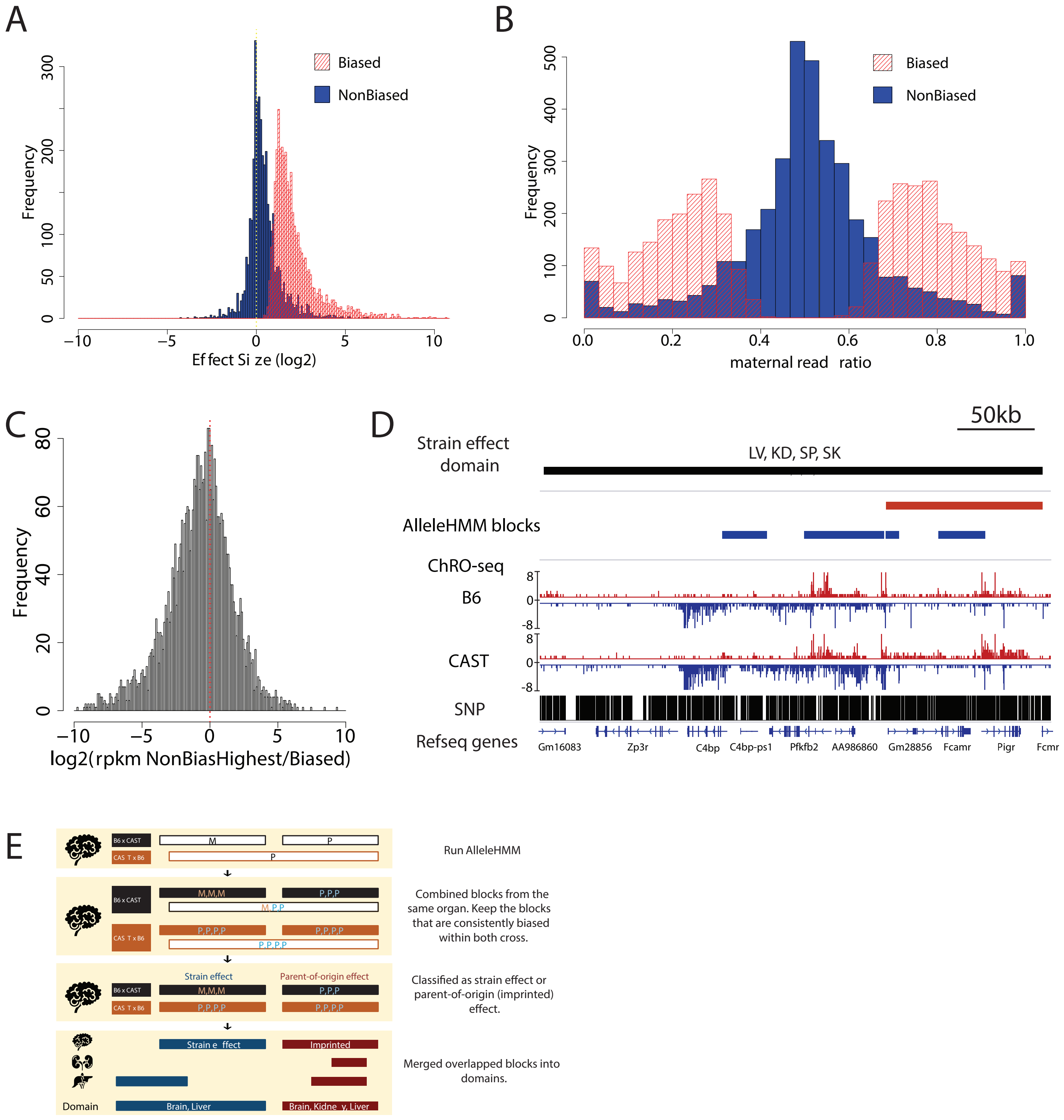


### Figure 1- figure supplement 1

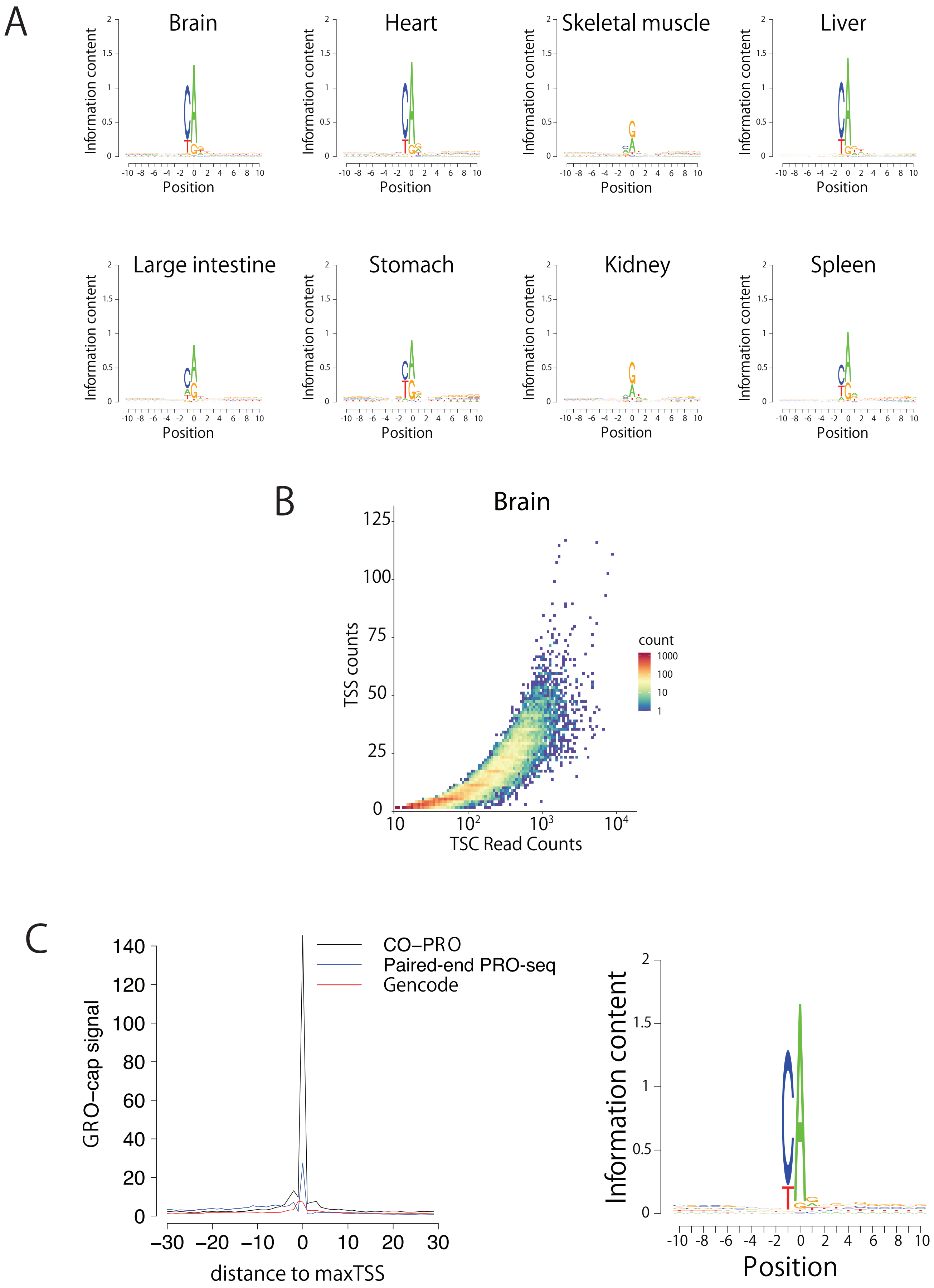


Figure 2 - figure supplement 1

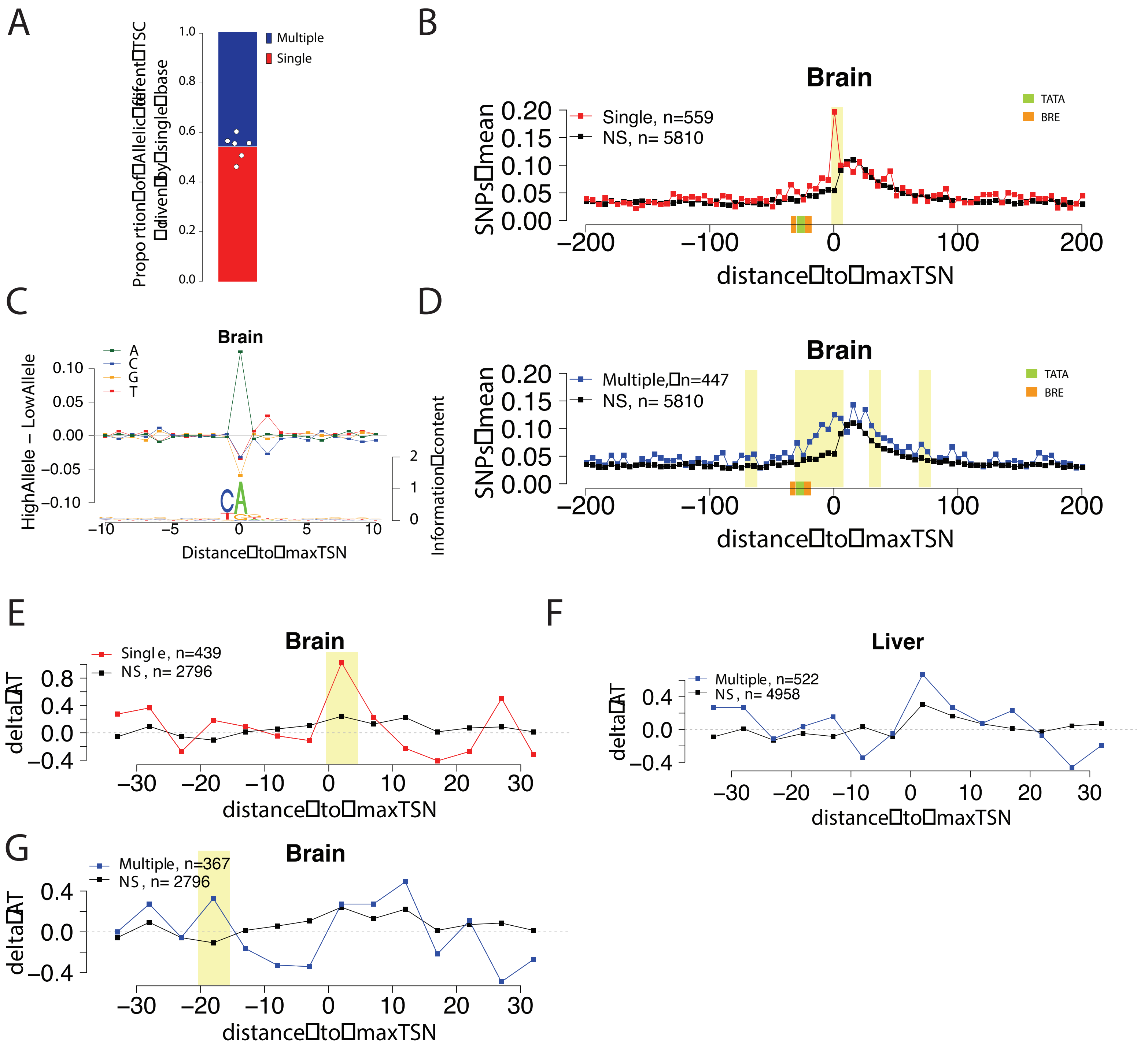


Figure 3 -figure supplement 1

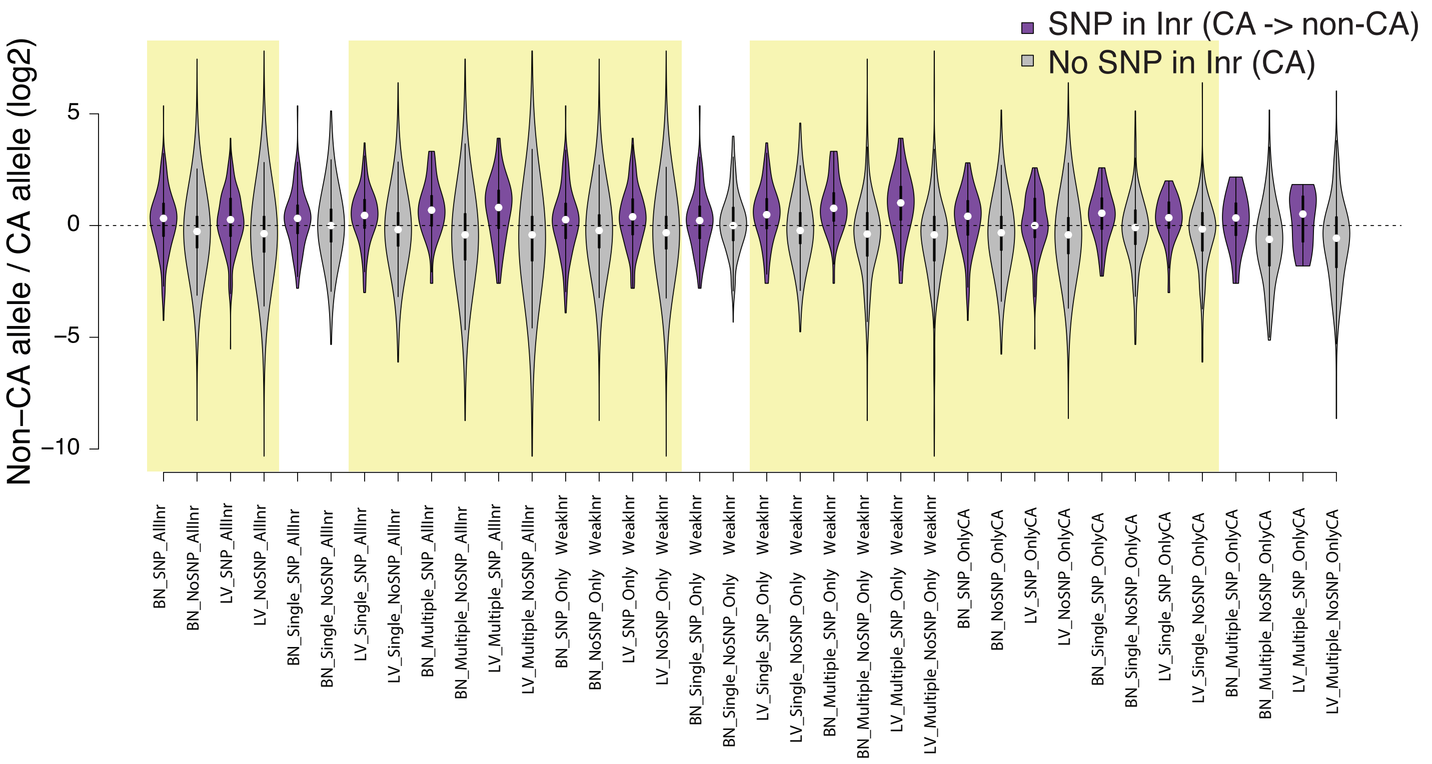


Figure 4 - figure supplement 1

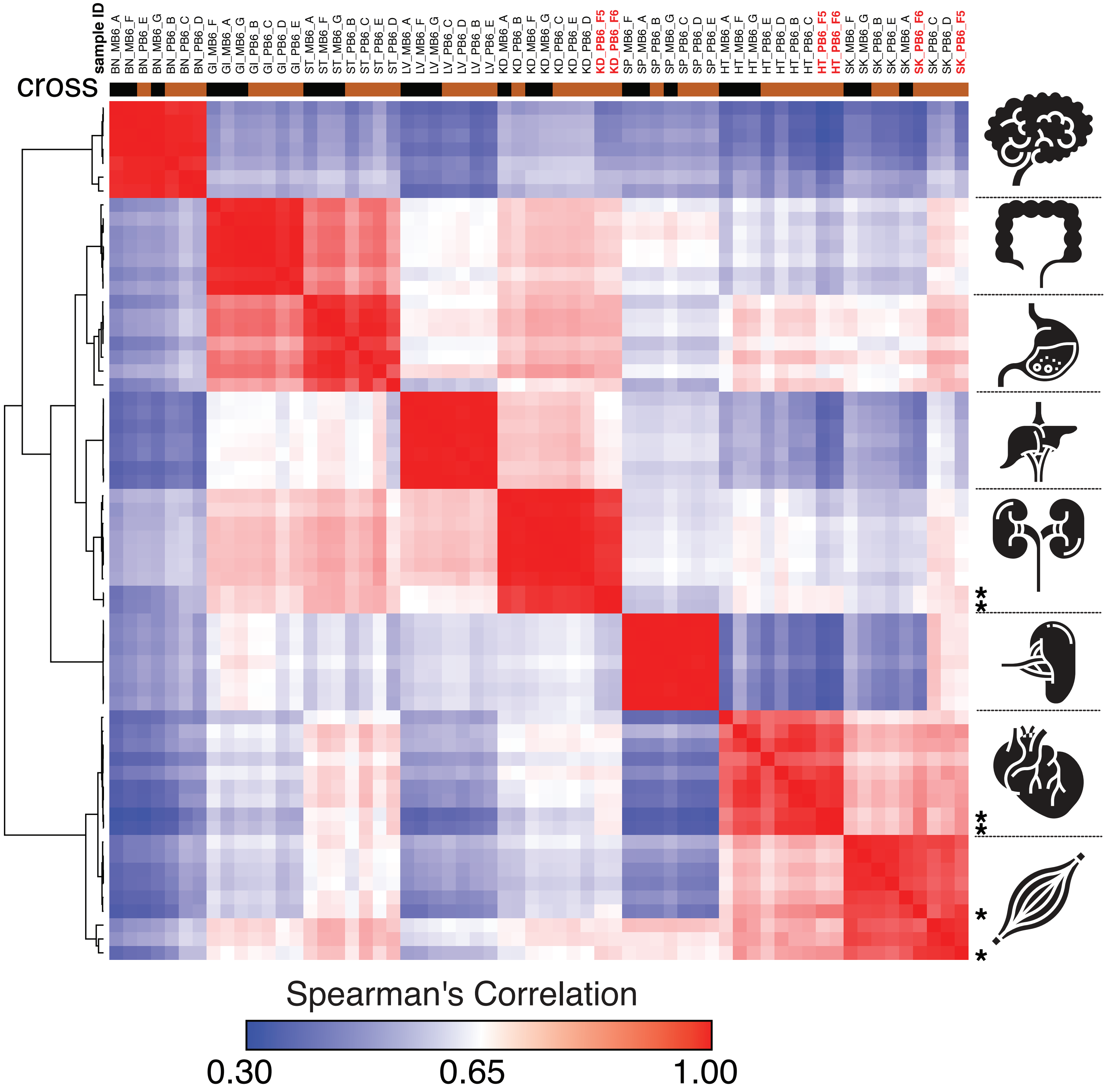


### Figure 5 - figure supplement 1

Spearman's rank correlation of the ChRO-seq data, including samples with a single base resolution for the Pol II active site. Samples with single nucleotide precision are shown on the top in red bold font and at the right with star. (BN:brain, GI: large intestine, ST: Stomach, LV:liver, KD: kidney, SP:spleen, HT: heart, SK:skeletal muscle, MB6: B6 x CAST, PB6: CAST x B6)

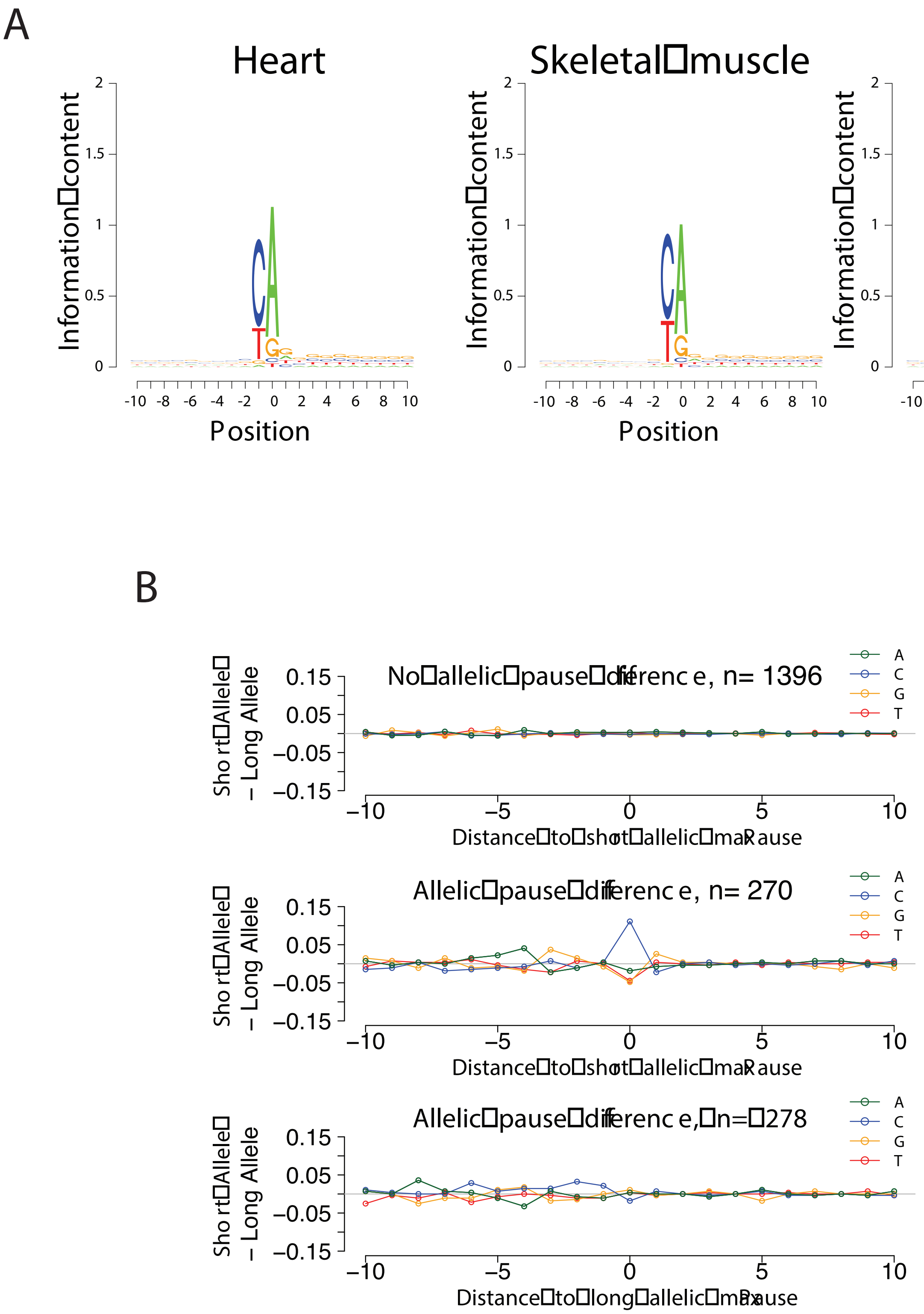


### Figure 5 - figure supplement 2

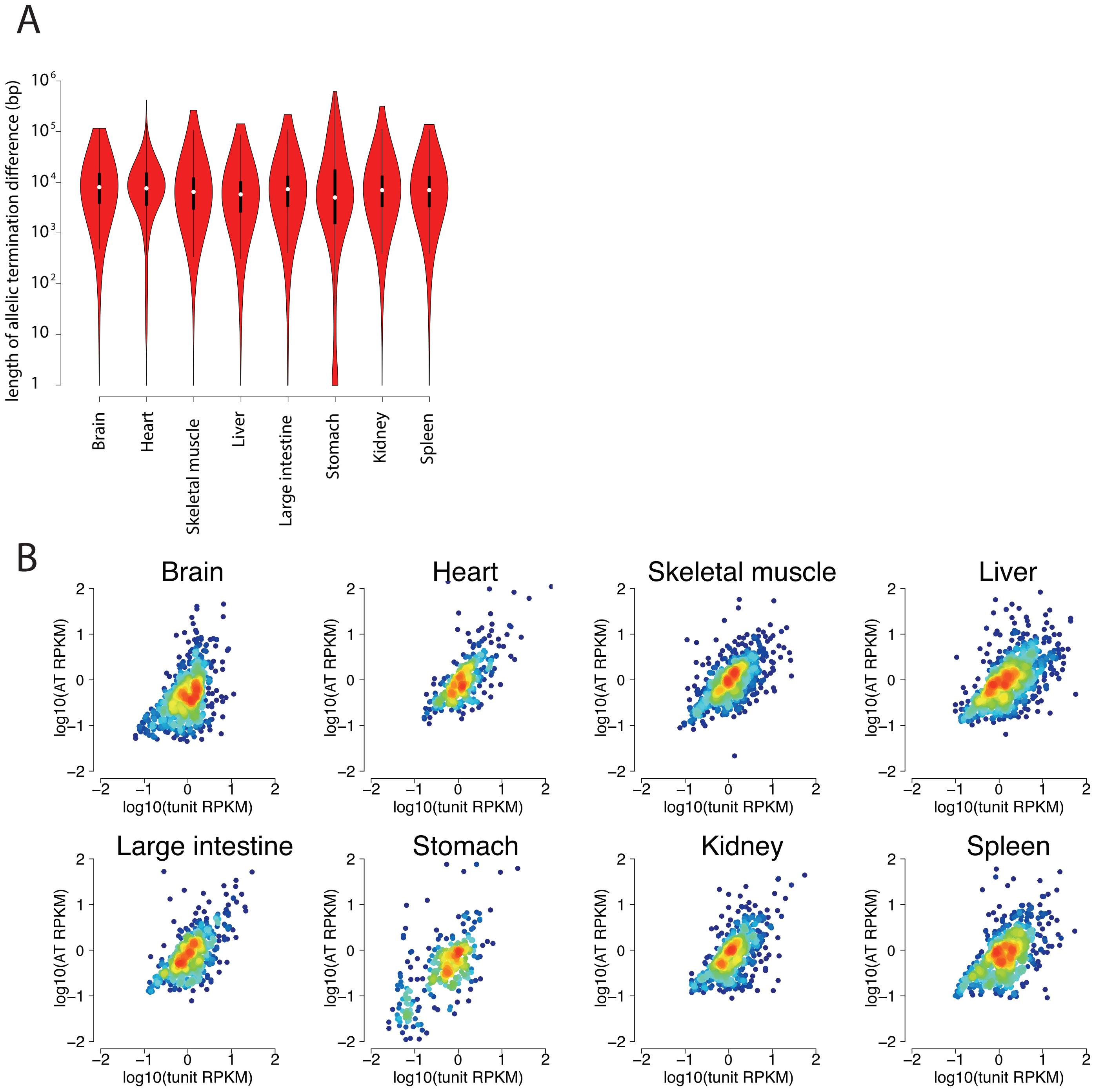


Figure 6 - figure supplement 1

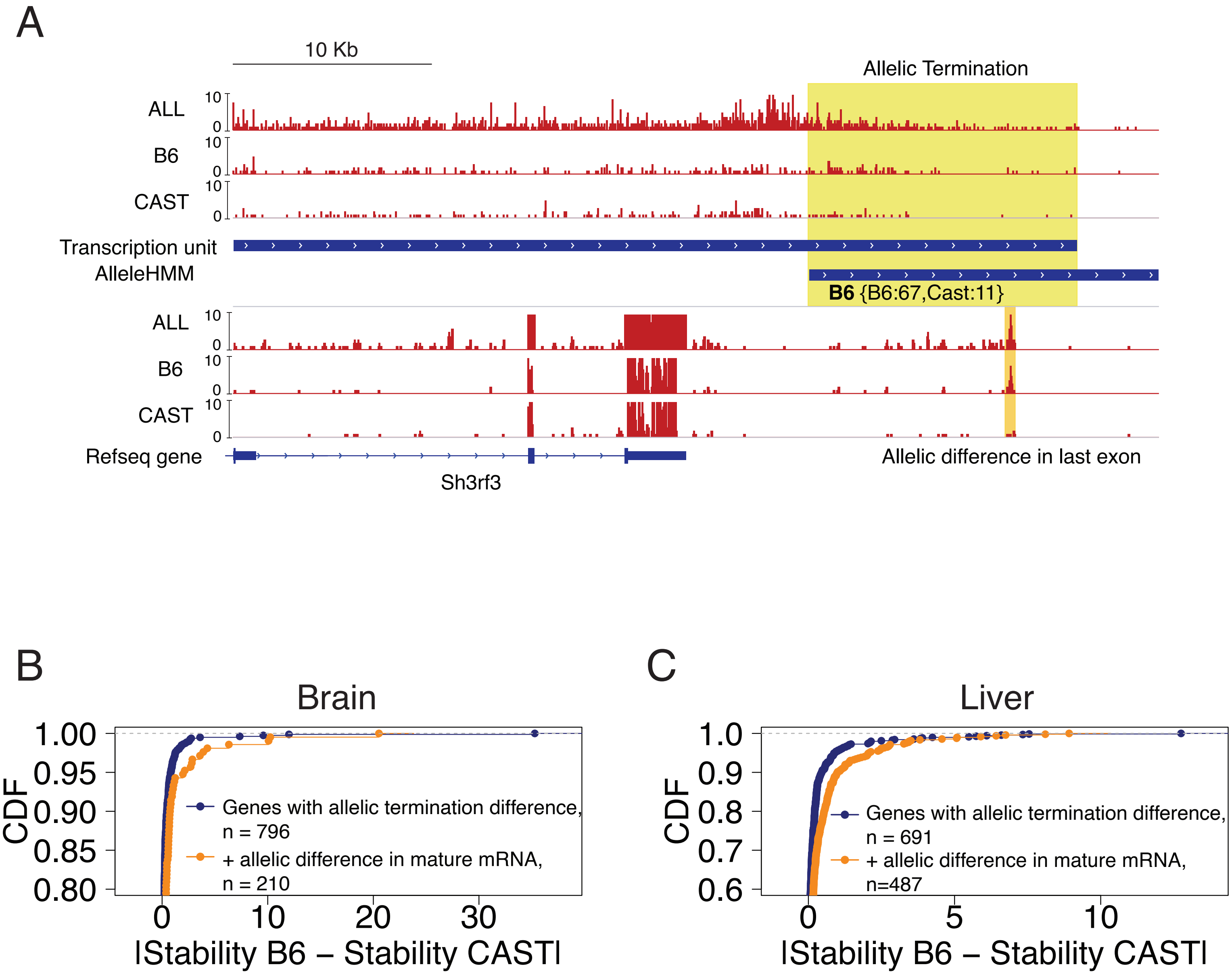


Figure 6 - figure supplement 2
